## Supporting information for "Potentiating antibiotic treatment using fitness-neutral gene expression perturbations"

**Affiliations:**
[1]Department of Chemical and Biological Engineering, University of Colorado at Boulder, Boulder, CO 80303, USA.
[2]Joint BioEnergy Institute, Emeryville, CA, 94608, USA.
[3]Biomass Science and Conversion Technologies, Sandia National Laboratories, Livermore, CA, 94550, USA.
[4]Theoretical Biology & Biophysics Group, Theoretical Division, Los Alamos National Laboratory, Los Alamos, NM 87545
[5]BioFrontiers Institute, University of Colorado at Boulder, Boulder, CO 80303, USA.

**This PDF file includes:**
Supplementary Figures 1-9
Supplementary Tables 1-2

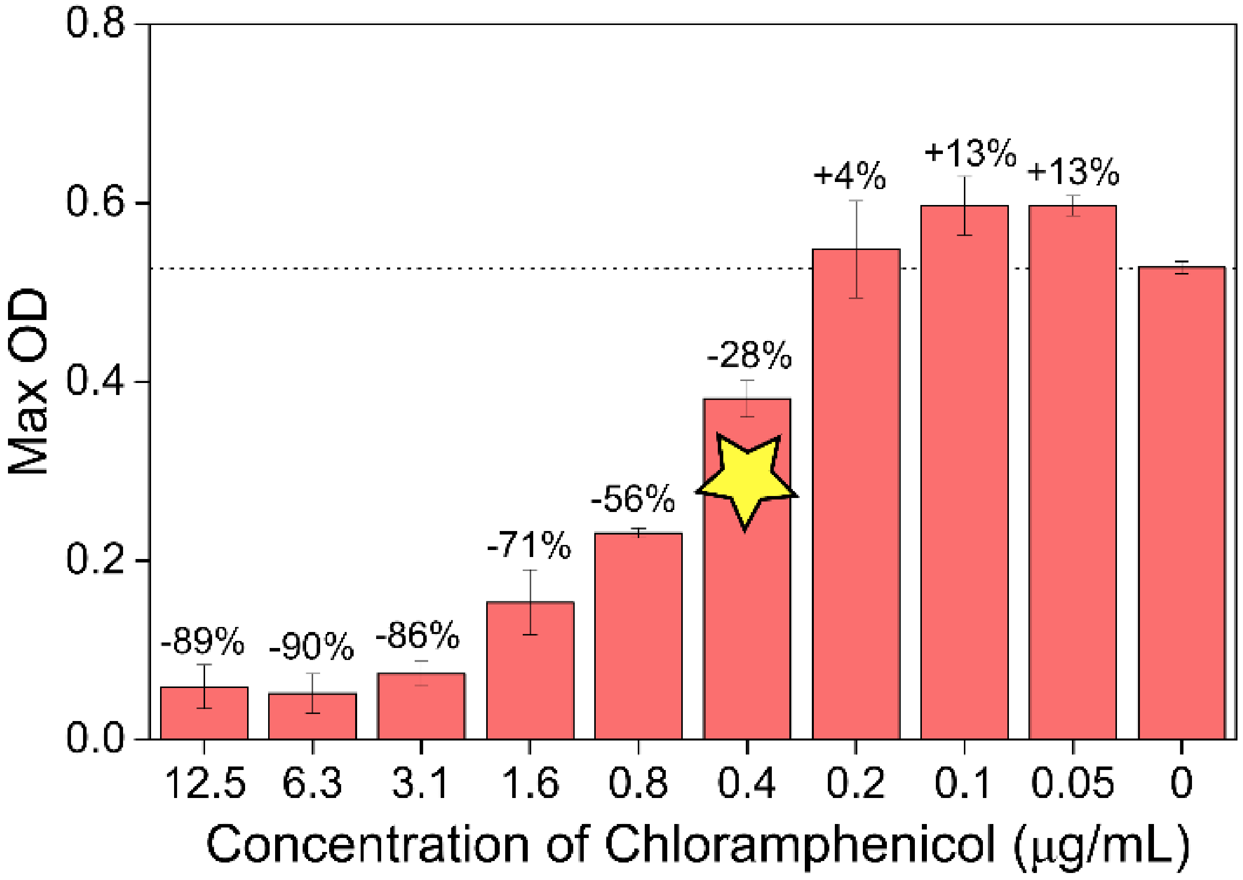


**Supplementary Figure 1.** Example of an antibiotic dilution test to identify the drug concentration suitable for combination therapy. E. coli BW25113 was grown in a range of concentrations for 16 hours. The concentration that resulted in a maximum optical density 10-50% lower than the no treatment case was selected for each antibiotic (indicated here with a star at 0.4 µg/mL of chloramphenicol). Error bars are SD, n = 3.

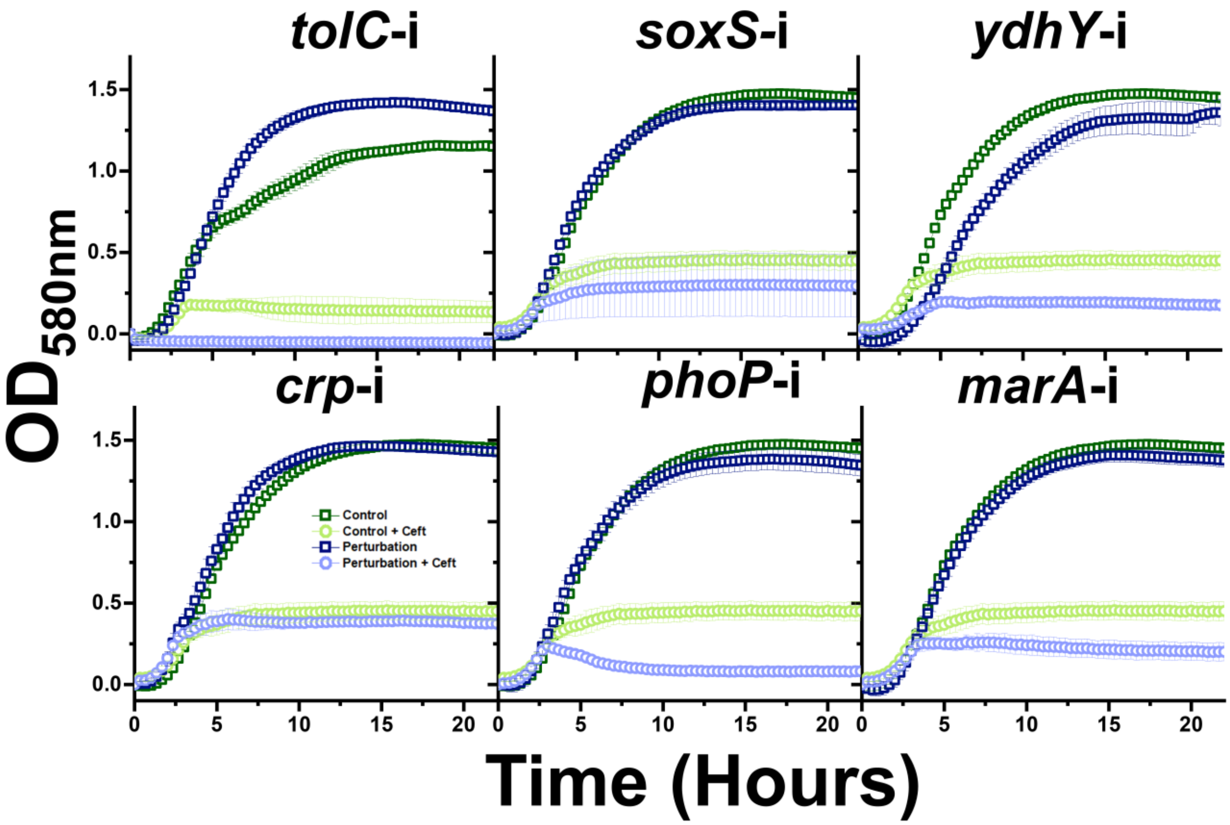


**Supplementary Figure 2.** Growth of CRISPRi strains during exposure to 2.0 µg/mL ceftriaxone in LB medium. Error bars represent standard deviation of four biological replicates. Growth is normalized to starting ODs.

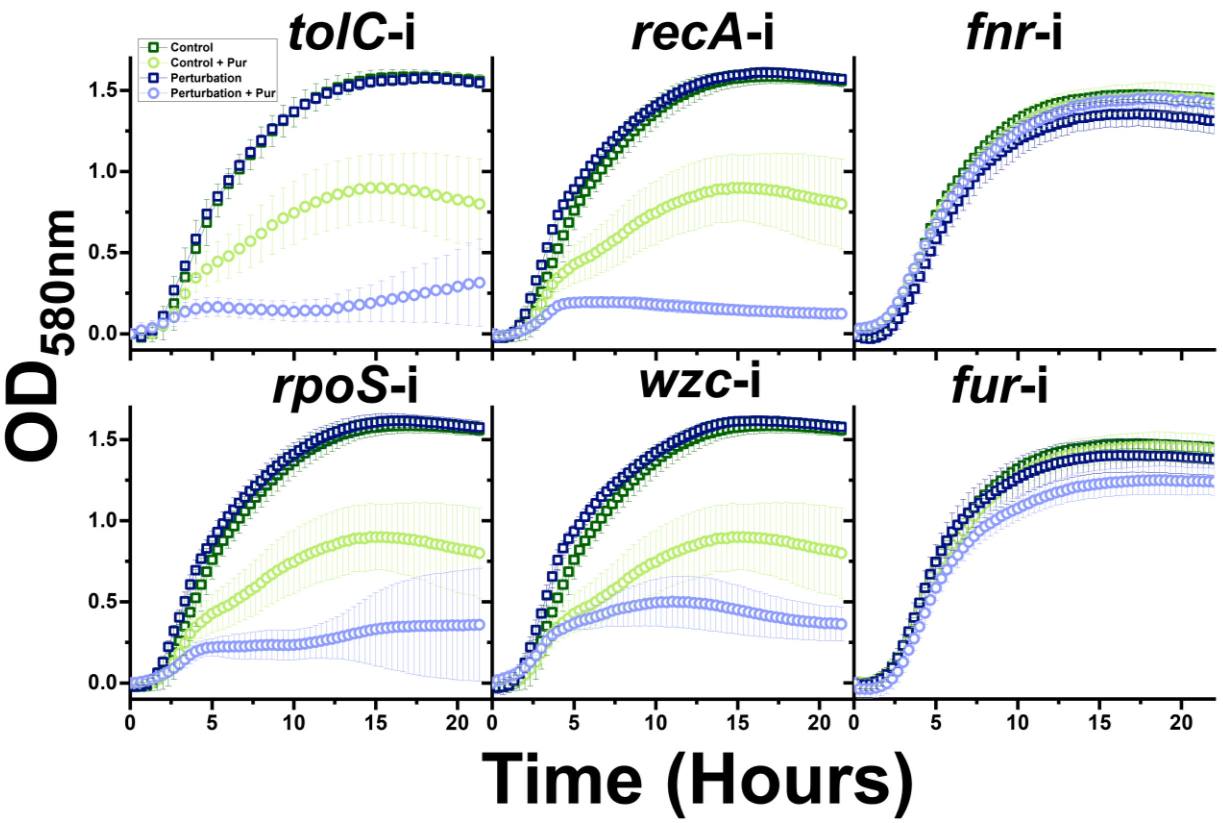


**Supplementary Figure 3.** Growth of CRISPRi strains during exposure to 50.0 µg/mL puromycin in LB medium. Error bars represent standard deviation of four biological replicates. Growth is normalized to starting ODs.

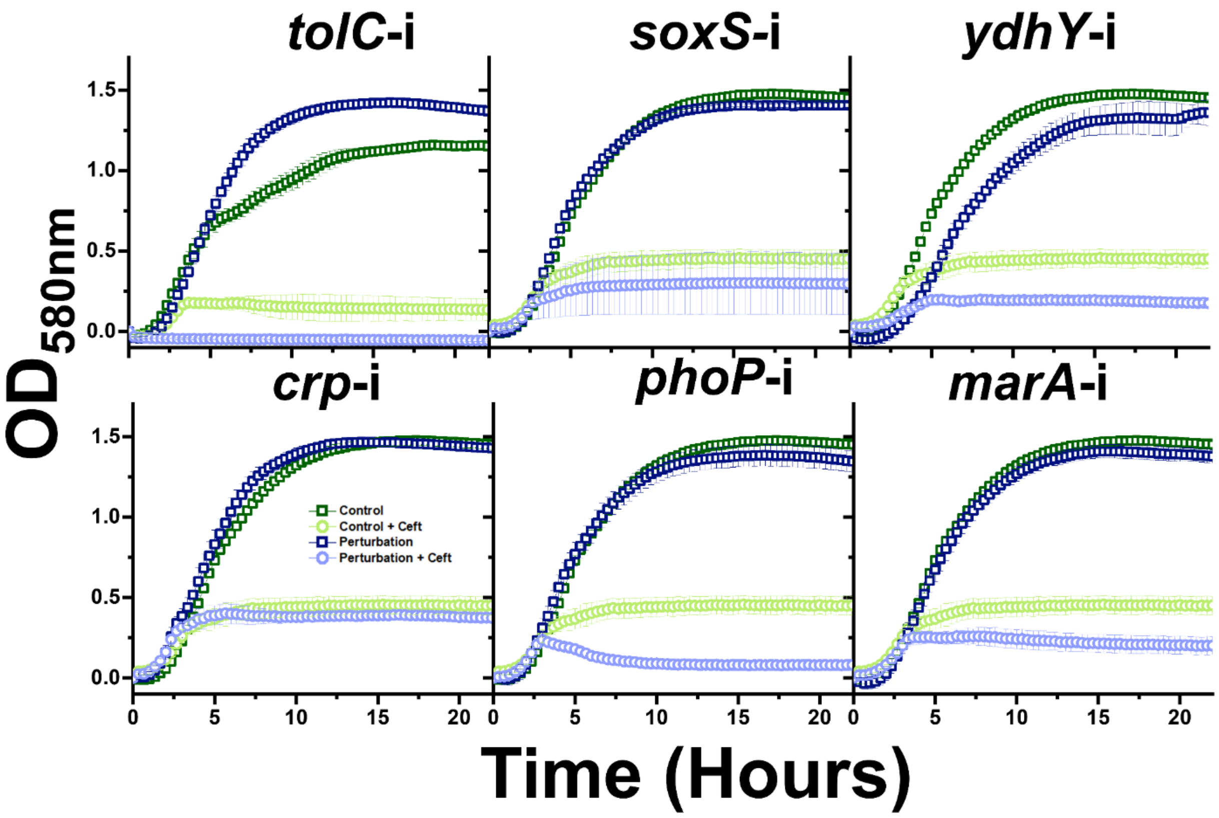


**Supplementary Figure 4.** Growth of CRISPRi strains during exposure to 0.25 µg/mL tetracycline in LB medium. Error bars represent standard deviation of four biological replicates. Growth is normalized to starting ODs.

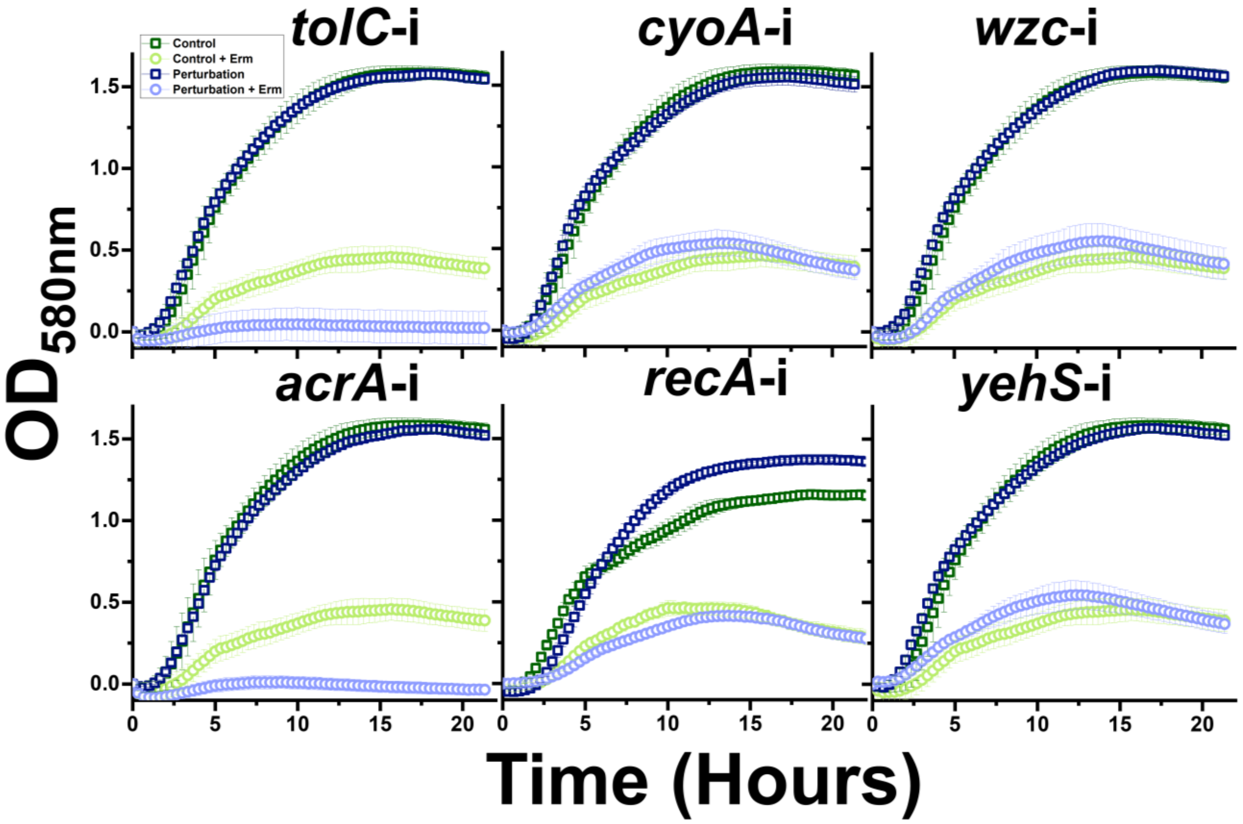


**Supplementary Figure 5.** Growth of CRISPRi strains during exposure to 50.0 μg/mL erythromycin in LB medium. Error bars represent standard deviation of four biological replicates. Growth is normalized to starting ODs.

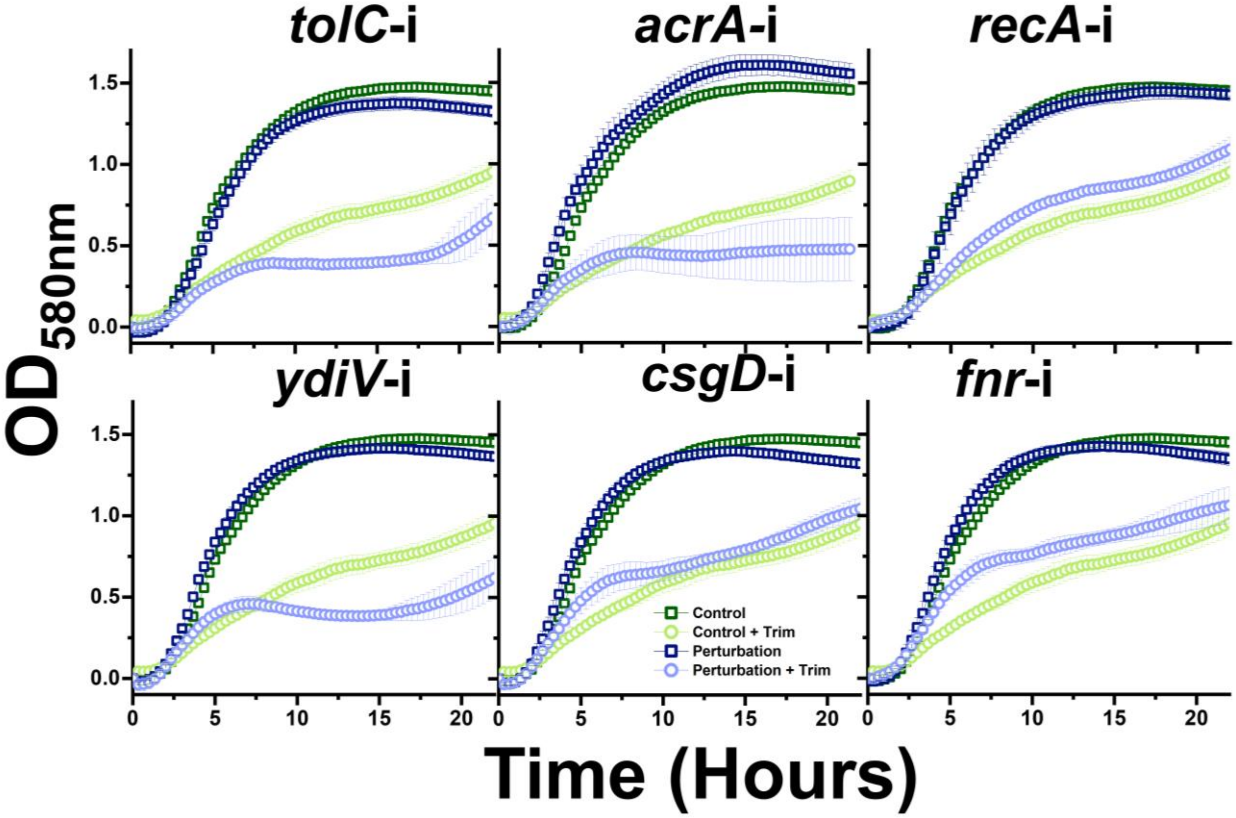


**Supplementary Figure 6.** Growth of CRISPRi strains during exposure to 0.125 µg/mL trimethoprim in LB medium. Error bars represent standard deviation of four biological replicates. Growth is normalized to starting ODs.

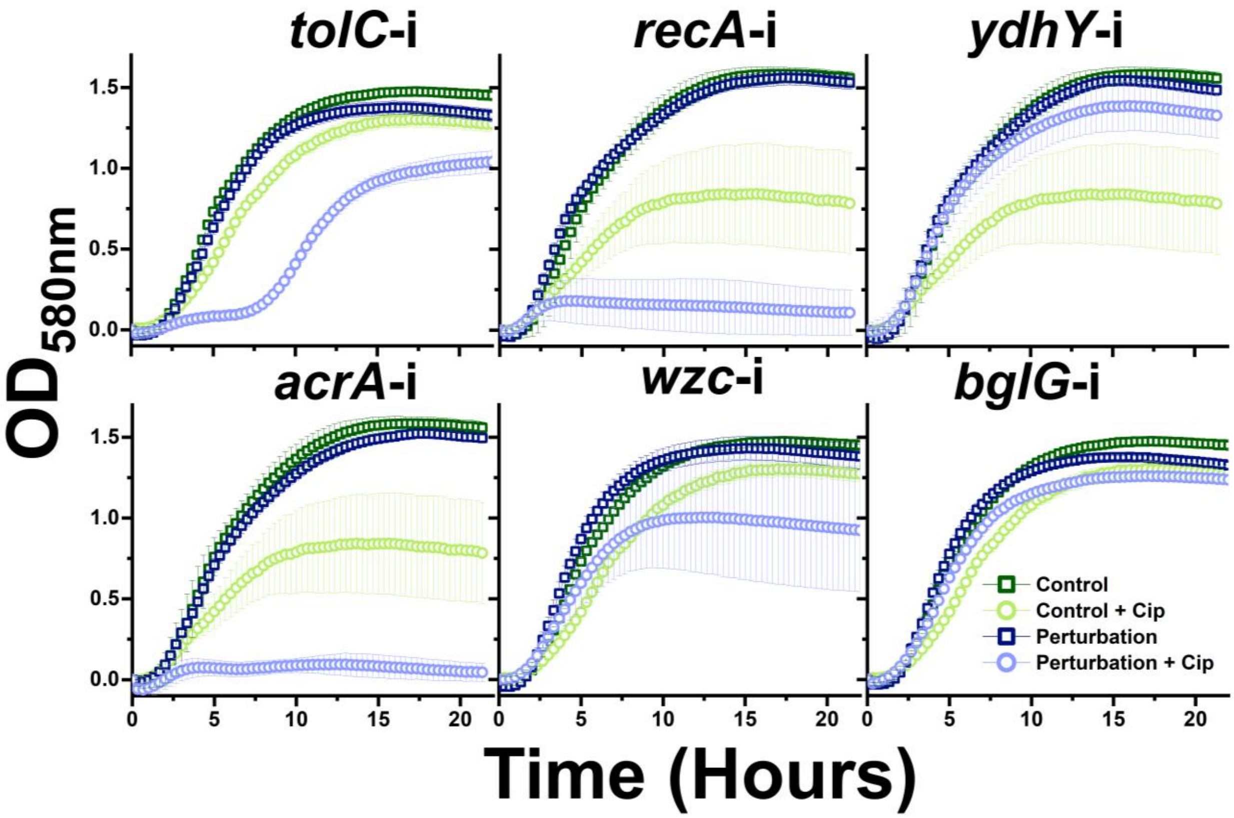


**Supplementary Figure 7.** Growth of CRISPRi strains during exposure to 0.008 μg/mL ciprofloxacin in LB medium. Error bars represent standard deviation of four biological replicates. Growth is normalized to starting ODs.

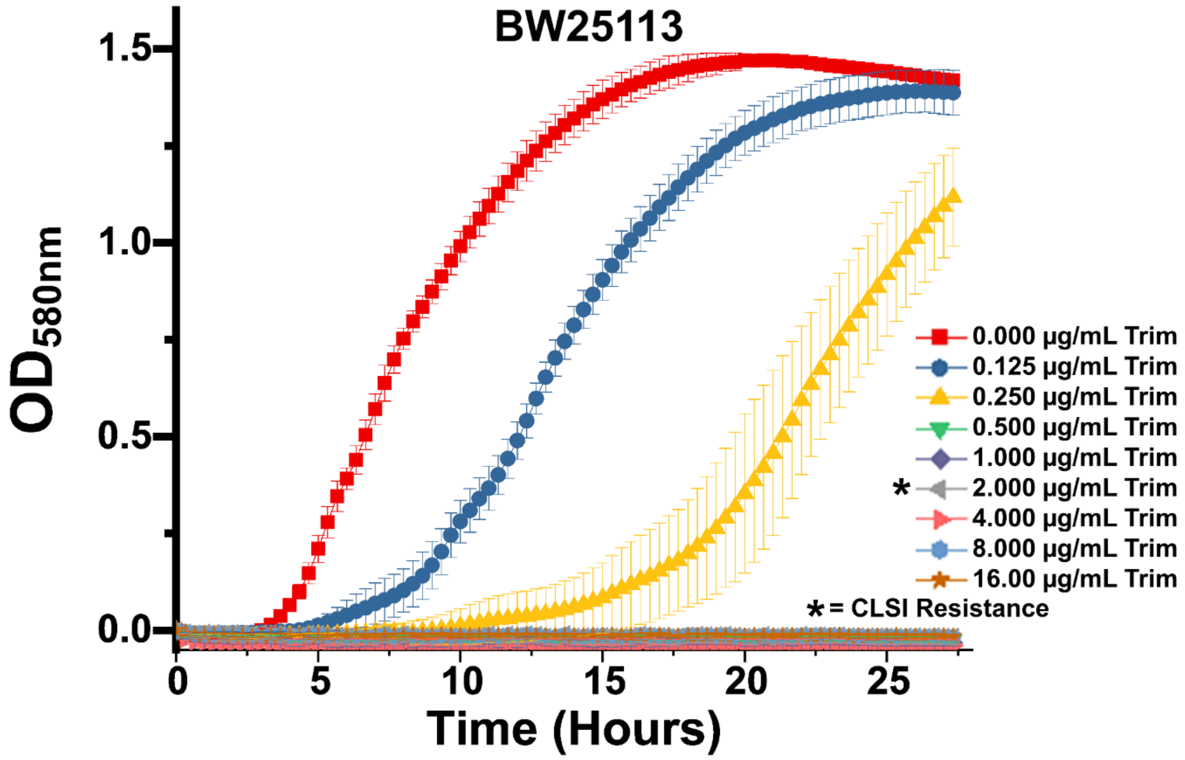


**Supplementary Figure 8.** BW25113 growth tests of trimethoprim resistance Cultures of BW25113 were grown in caMHB for 24 hours to quantify basal *E. coli* resistance to trimethoprim. Cells were unable to survive 0.5 µg/mL trimethoprim and above, 4-fold below the CLSI breakpoint for trimethoprim resistance. Error bars represent standard deviation of biological triplicates.

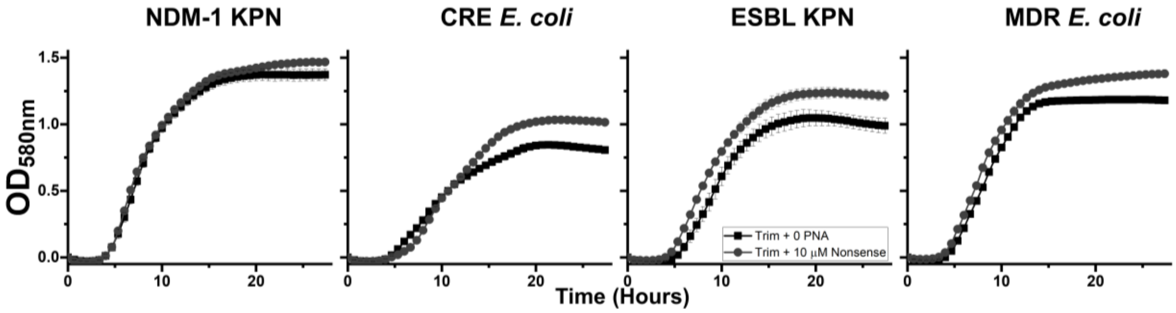


Supplementary Figure 9. Growth of clinically isolated MDR bacteria during exposure to 2.0 μg/mL trimethoprim in the presence or absence of 10 μM nonsense targeting PNA. Error bars represent standard deviation of biological triplicates.

Table S1. sgRNA targets examined in this study.

| Target | Sequence (5' to 3') |
|---|---|
| soxS | ctacatcaatgttaagcggc |
| tolC | ggctcaggccgataagaatg |
| acrA | agcatcagaacgaccgccag |
| ydhY | gatcgtccactattagatat |
| crp | aaacagacccgactctcgaa |
| phoP | agaaataaaaatgcgcgtac |
| marA | ccagtccaaaatgctatgaa |
| recA | taccaaattgtttctcaatc |
| wzc | caacatgccgctccggtaac |
| bglG | aattctcaacaataatgttg |
| rpoS | actgggttcctgttctacta |
| yehS | gcgtgcgctacattttgaaa |
| csgD | aatgaagtccatagtattca |
| fnr | tcccggaaaagcgaattata |
| fur | aataccgccctaaagaaagc |
| cyoA | aggaaatacaataaaagttt |
| ydiV | aatcagaatgataaagattc |
| rfp (nonsense control) | aactttcagtttagcggtct |

Table S2. PNA targets examined in this study.

| Target | Sequence (N to C terminus) |
|---|---|
| AcrA | KFFKFFKFFK-O-tatgtaaacctc |
| Fnr | KFFKFFKFFK-O-gatcataggtct |
| CsgD | KFFKFFKFFK-O-tgatgaaacccc |
| RecA | KFFKFFKFFK-O-gtcgatagccat |
| Nonsense | KFFKFFKFFK-O-gaataagggcga |
